# Rats emit unique distress calls in social inequality conditions

**DOI:** 10.1101/2022.02.22.481162

**Authors:** Shota Okabe, Yuki Takayanagi, Masahide Yoshida, Tatsushi Onaka

## Abstract

Humans show aversion toward inequality of social reward, and this aversion plays important roles for the establishment of social cooperation. However, it has remained unknown whether commonly used experimental animals show negative responses to social reward inequality. In this study, we found that rats showed bonding-like behavior to an affiliative human who repeatedly stroked the rats. In addition, these rats emitted distress calls, an index of negative emotion, when an affiliative human stroked another rat in front of them. These distress calls had acoustic characteristics different from those emitted in response to physical stress stimuli such as air-puff. Rats emitted calls with higher frequency (28 kHz) and shorter durations (0.05 sec) in an inequality condition than the frequency and durations of calls emitted when receiving air-puff. Our results suggested that rats exhibited negative emotion with unique distress calls in response to a social inequality condition.

## Introduction

It is well known that humans show negative responses in a situation of reward inequality. A sense of fairness and expression of disgust-related negative emotion toward reward inequality, which are considered to facilitate collaboration among cooperative group members, especially in a hunting and gathering society, have been proposed to have evolved during the course of evolution ^1,2^. Recent research suggests that not only primates but also dogs^3,4^ and crows^5^ exhibit negative responses in inequality conditions. In addition, some studies have revealed that rodents^6,7^ also exhibit negative responses in inequality conditions. These animal studies suggested that a sense of fairness is not specific to primates. In the majority of experiments using animals for investigation of reward inequality, food has been used as a resource of reward to be distributed. On the other hand, humans show negative responses not only toward inequality of money or food reward but also toward inequality of social relationship-related reward. For example, humans feel negative emotions, envy or jealousy, in social situations. However, little is known about the neural mechanisms underlying responses to social reward inequality. It was not known whether animals show negative emotion toward an inequitable distribution of social rewards.

In our previous study, we found that rats that received repeatedly stroked stimuli during the adolescent period show affiliative responses toward humans. These stroked rats also showed preference for interacting with humans. For these stroked rats, stroking stimuli had a reward value, which induces conditioned place preference ^8,9^. Humans feel a negative emotion, jealousy, when affiliative specific persons show an affiliative attitude toward others instead of them. If rats can form an affiliative relationship towards a specific human, it may be possible to examine whether rats show a jealousy-like response, although it has remained to be determined whether rats show affiliative responses selectively towards specific humans.

Rats emit two broad subgroups of ultrasonic vocalizations (USVs) depending on their emotional states; one is a USV with a frequency range of 18-32 kHz, referred to as “22-kHz calls”^10,11^, which are emitted in aversive states such as exposure to predators. The other is a USV with a frequency of more than 35 kHz, referred to as “50-kHz calls” ^11^, which are emitted in appetitive or rewarding situations such as social play. By analyzing the USVs emitted in inequality conditions, we could estimate whether rats show negative or appetitive emotion.

The aim of this study was to determine whether rats show negative emotion toward a social inequality condition. To achieve this aim, we first examined whether rats form an affiliative relationship with a specific human. We then recorded USVs to reveal whether rats show negative emotion in a situation when the affiliative human stroked another rat. Oxytocin is known to play an important role in social relationships. In previous studies, we revealed that the activity of oxytocin neurons and affiliative behavior toward a human were correlated^8^. Thus, we tested whether inhibition of oxytocin function would disrupt affiliative relationships between rats and humans and prevent rats from showing negative responses in social inequality conditions.

## Results

### Post-weaning repeated affiliative touches result in the development of bonding between rats and humans

To investigate whether rats emit distress calls during an inequality social situation, we initially produced affiliative rats by giving gentle touches of stroking stimuli or tickling during the post-weaning period according to procedures reported previously ^8^ with minor revisions. In the developmental period, we recorded USVs during stroking stimuli every week (Fig. 1a) and counted the numbers of 50-kHz and 22-kHz calls (Fig. 1b, c) to confirm affiliative responsiveness. Repeated measures one-way ANOVA revealed a significant effect of postnatal weeks of age [F (8, 88) = 15.608, *P* < 0.001] in the numbers of 50-kHz calls. The numbers of 50-kHz calls at the ages of 5, 6, 7, 8, 9, 10, and 11 weeks were significantly increased compared to the number of calls at 3 weeks of age (3 weeks versus 5 weeks, *P* = 0.026; 3 weeks versus 6 weeks, *P* = 0.015; 3 weeks versus 7 weeks, *P* = 0.013; 3 weeks versus 8 weeks, *P* = 0.019; 3 weeks versus 9 weeks, *P* = 0.006; 3 weeks versus 10 weeks, *P* = 0.014; 3 weeks versus 11 weeks, *P* = 0.02) (Fig. 1d). The numbers of 50-kHz calls at the ages of 7, 8, 9, 10, and 11 weeks were significantly increased compared to the number of calls at 4 weeks of age (4 weeks versus 7 weeks, *P* = 0.023, 4 weeks versus 8 weeks, *P* = 0.048; 4 weeks versus 9 weeks, *P* = 0.009; 4 weeks versus 10 weeks, *P* = 0.039; 4 weeks versus 11 weeks, *P* = 0.029). In addition, the number of 50-kHz calls at the age of 9 weeks was significantly increased compared to the numbers of calls at 5 and 6 weeks of age (5 weeks versus 9 weeks, *P* = 0.02; 6 weeks versus 9 weeks, *P* = 0.027). The number of 22-kHz calls during stroking was small and did not significantly change during the developmental period (Fig. 1e). These data show that rats that received stroking stimuli during the post-weaning period showed an affiliative response toward stroking stimuli.

**Fig. 1:**
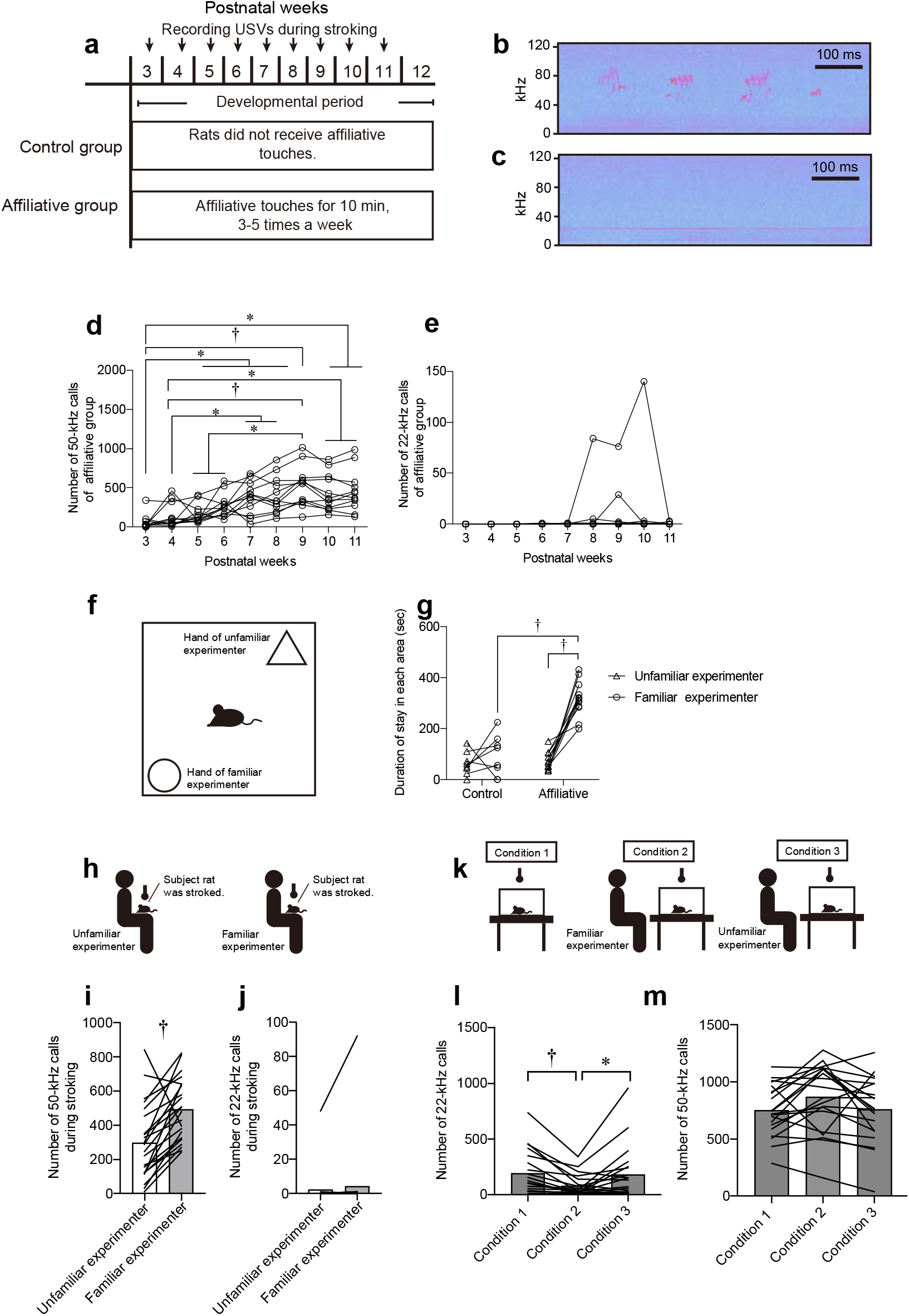
Post-weaning repeated affiliative touches result in bonding between rats and humans. Time schedules of experiments. Ultrasonic vocalizations were recorded for 5 min once per week between 3 and 11 weeks of age (developmental period) in the affiliative touch group (a). Spectrograms of 50-kHz (b) and 22-kHz calls (c). Time courses of the numbers of vocalizations during stroking (d, 50-kHz; e, 22-kHz). (f) Illustration of a social preference test. (g) Durations spent staying in the familiar experimenter’s hand area compared with durations spent staying in the unfamiliar experimenter’s hand area in a social preference test. (h) Illustrations of experimental procedures. USVs were recorded when rats were stroked by an unfamiliar or familiar experimenter. Numbers of 50-kHz (i) and 22-kHz (j) calls. (k) Illustrations of experimental procedures. USVs were recorded in the isolation condition (condition 1). USVs were also recorded when a familiar experimenter (condition 2) or unfamiliar experimenter (condition 3) sat down in front of the observer’s cage. (l, m) Numbers of 22-kHz calls (l) and 50-kHz calls (m). †, *P* < 0.01, *, *P* < 0.05.

Next, we examined whether rats show an affiliative response to a specific experimenter (Fig. 1f). Repeated measures two-way ANOVA revealed significant effects of group [F (1, 18) = 50.609, *P* < 0.001] and experimenter [F (1, 18) = 40.832, *P* < 0.001] and a significant interaction of the two [F (1, 18) = 24.827, *P* < 0.001] in times spent in hand areas in a preference test. The time spent staying in the familiar experimenter’s hand area was significantly longer than that in the unfamiliar experimenter’s hand area in the affiliative group (*P* < 0.001, post-hoc Holm’s test). The duration of staying in the familiar experimenter’s hand area in the affiliative group was also significantly longer than that in the control group (*P* < 0.001, post-hoc Holm’s test) (Fig.1g). Rats in the non-stroked control group did not show any preference (Fig. 1g). These data indicate that rats in the affiliative group showed preference toward a familiar experimenter.

We counted the numbers of USVs during stroking stimuli that were given by a familiar experimenter and by an unfamiliar experimenter (Fig.1h). The number of 50-kHz calls was significantly larger during stroking stimuli given by a familiar experimenter than that during stroking stimuli given by an unfamiliar experimenter (*P* = 0.001, Wilcoxon signed-rank test) (Fig.1i). The number of 22-kHz calls did not significantly differ between the conditions of stroking stimuli by familiar and unfamiliar experimenters (Fig.1j).

Social isolation in an unfamiliar environment induces stress responses in social animals ^12^, and being accompanied by affiliative individuals attenuates the isolation-induced stress responses ^12,13^. In order to determine whether rats showed negative emotion during isolation and whether isolation-induced negative emotion was abolished by the presence of a familiar experimenter, we examined 22-kHz USV as an index of stress responses in an isolation condition and in the condition of isolation with the presence of a human (Fig. 1k). In a condition of isolation (condition 1), a subject rat was isolated in a clean cage. In a condition of isolation in the presence of a familiar experimenter (condition 2), a subject rat was kept in a clean cage and a familiar experimenter, who gave affiliative touches during the developmental period, sat down in front of the cage. In another condition of isolation in the presence of an unfamiliar experimenter (condition 3), a subject rat was kept in a clean cage and an unfamiliar experimenter sat down in front of the cage. In the numbers of 22-kHz calls, repeated measures one-way ANOVA revealed a significant effect of condition [F (2, 38) = 5.747, *P* = 0.009]. The number of 22-kHz calls in the presence of a familiar experimenter (condition 2) was significantly smaller than that in the social isolation condition (condition 1) or in the presence of an unfamiliar experimenter (condition 3) (condition 1 versus condition 2, *P* = 0.002; condition 3 versus condition 2, *P* = 0.029, post-hoc Holm’s test) (Fig. 1l). The significant reduction in the number of 22-kHz calls in the presence of a familiar experimenter indicates that the presence of a familiar experimenter suppressed isolation-induced negative emotion in rats. In the number of 50-kHz calls, repeated measures one-way ANOVA revealed a significant effect of condition [F (2, 38) = 3.589, *P* = 0.037]. However, there were no significant differences in the number of 50-kHz calls among these conditions (post-hoc Holm’s test, Fig. 1m). These results showed that rats could discriminate between a familiar experimenter and an unfamiliar experimenter and that repeated affiliative touches during the adolescent period resulted in the formation of a bonding between rats and an familiar experimenter.

### Affiliative rats emitted 22-kHz calls during a social inequality condition

In order to determine whether rats showed negative emotion in a social inequality situation, we also examined 22-kHz USVs in inequality conditions. Rats were placed in a social isolation condition (condition 1), in the presence of a familiar experimenter (condition 2), or in an inequality condition of condition 4 or condition 5 (Fig. 2a). A subject rat (observer) was kept in a clean cage and a familiar experimenter stroked its cage mate (condition 4) or an unfamiliar rat (condition 5) in front of the cage.

**Fig. 2:**
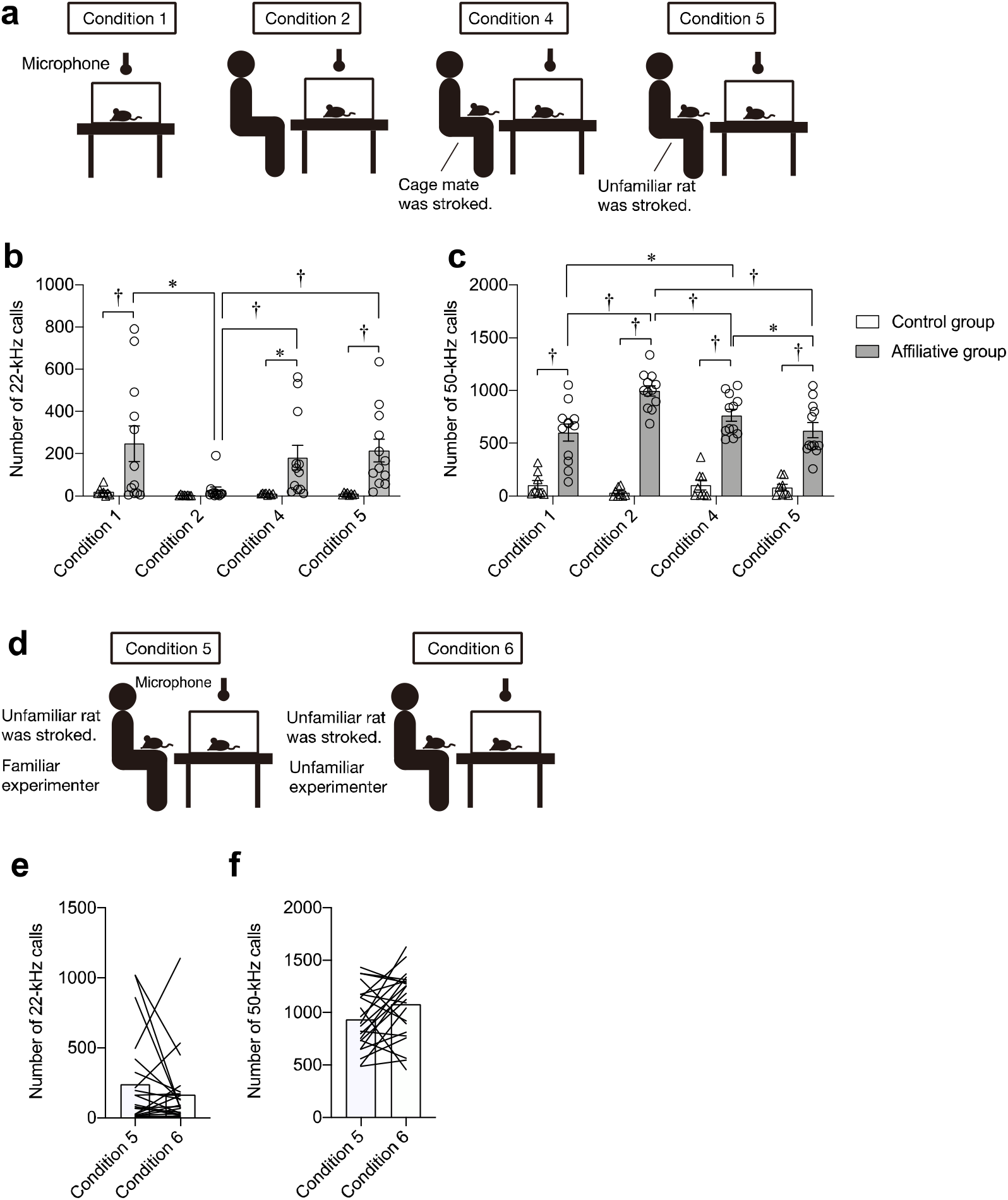
Affiliative rats emitted 22-kHz calls during the inequality condition. (a) Illustrations of experimental procedures. USVs were recorded in four conditions. In condition 1, subject rats were isolated in a clean cage. In condition 2, subject rats were kept in a clean cage and a familiar experimenter who had stroked the rats during the developmental period sat down in front of that cage. In condition 4, subject rats (observers) were kept in a clean cage and a familiar experimenter stroked cage mates (demonstrators) of subject rats in front of that cage. In condition 5, rats were kept in a clean cage and a familiar experimenter stroked unfamiliar rats (demonstrators) in front of that cage. (b) Numbers of 22-kHz calls during conditions 1, 2, 4, and 5. (c) Numbers of 50-kHz calls during conditions 1, 2, 4, and 5. (d) Illustrations of experimental procedures. The observer rat’s USVs were recorded when a familiar experimenter (condition 5) or unfamiliar experimenter (condition 6) stroked the demonstrator rat in front of the observer’s cage. Numbers of 22-kHz (e) and 50-kHz (f) calls. †, *P* < 0.01, *, *P* < 0.05. Error bars denote standard error of the mean.

In the numbers of 22-kHz calls, repeated measures two-way ANOVA revealed significant effects of group [F (1, 18) = 7.699, *P* = 0.012] and condition [F (3, 54) = 4.116, *P* = 0.023] and a significant interaction of the two [F (3. 54) = 3.258, *P* = 0.049]. The number of 22-kHz calls in the affiliative group was significantly larger than the number of 22-kHz calls in the non-stroked control group in the condition of social isolation (condition 1, *P* = 0.002). The number of 22-kHz calls in the affiliative group was significantly smaller in the presence of a familiar experimenter (condition 2) than in a condition of social isolation (condition 1) (condition 2 versus condition 1, *P* = 0.011, post-hoc Holm’s test). The results showed that the isolation-induced increase in the number of 22 kHz calls was abolished by the presence of a familiar experimenter.

Interestingly, the numbers of 22-kHz calls in affiliative groups were increased when the familiar experimenter stroked demonstrator familiar rats or unfamiliar rats in front of observer rats (conditions 4 and 5) compared with the number of 22-kHz calls in non-stroked control groups (Fig. 2b, in condition 4, *P* = 0.02; in condition 5, *P* = 0.006, post-hoc Holm’s test; Supplementary Movie 1 and 2). The numbers of 22-kHz calls in affiliative groups when the experimenter stroked demonstrator rats (conditions 4 and 5) were also significantly larger than the number of 22-kHz calls when the experimenter just sat in front of the observer rats (condition 2) (Fig. 2b, condition 2 versus condition 4, *P* = 0.005; condition 2 versus condition 5, *P* = 0.001, post-hoc Holm’s test). These results suggested that rats in the affiliative group showed negative emotion in response to an inequality condition. Our results showing no significant difference between the numbers of 22-kHz calls in conditions 4 and 5 suggested that the familiarity between an observer rat and demonstrator rat did not play a crucial role in the negative reaction to an inequality condition.

In the number of 50-kHz calls, repeated measures two-way ANOVA revealed significant effects of group [F (1, 18) = 98.746, *P* < 0.001] and condition [F (3, 54) = 6.394, *P* = 0.001] and a significant interaction of the two [F (3. 54) = 11.751, *P* < 0.001]. The numbers of 50-kHz calls in the affiliative groups were significantly larger than those in non-stroked control groups in all conditions (in conditions 1, 2, 4, and 5, *P* < 0.001, post-hoc Holm’s test) (Fig. 2c). The number of 50-kHz calls in the presence of the affiliative experimenter in the affiliative group (condition 2) was significantly larger than the numbers of 50-kHz calls in the other three conditions (condition 2 versus condition 1, *P* < 0.001; condition 2 versus condition 4, *P* = 0.001; condition 2 versus condition 5, *P* < 0.001, post-hoc Holm’s test). The number of 50-kHz calls in the affiliative group in a condition in which the familiar experimenter stroked a cage mate (condition 4) was significantly larger than the numbers of 50-kHz calls in the condition with no presence of the familiar experimenter (condition 1) and in the condition in which the familiar experimenter stroked an unfamiliar rat (condition 5) (condition 4 versus condition 1, *P* = 0.019; condition 4 versus condition 5, *P* = 0.014, post-hoc Holm’s test). The number of 50-kHz calls in the presence of the familiar experimenter was significantly reduced when the experimenter stroked another rat, especially an unfamiliar rat, suggesting that the presence of novel demonstrator rats reduced positive emotions.

Envy and jealousy have long been regarded as distinct emotions ^14^. Envy occurs when a person recognizes superior quality and achievement of other individuals. In contrast to envy, jealousy occurs in the context of social relationships among three individuals. For example, a feeling of jealousy occurs when an affiliative familiar person directs his/her attention toward other persons but not to oneself. Thus, we also examined the effects of familiarity of experimenters on the emission of 22-kHz calls of affiliative rats in another series of experiments. A subject rat (observer) was kept in a clean cage and the familiar experimenter (condition 5) or unfamiliar experimenter (condition 6) stroked a demonstrator rat in front of the observer rat (Fig. 2d). There was no significant difference in the numbers of 22-kHz and 50-kHz calls of the observer rats between the condition 5 and condition 6 (Fig. 2e, f). The results suggested that the familiarity of the experimenter did not significantly affect the emission of calls in the inequality condition. Thus, it was possible that rats emitted 22-kHz calls not because of jealousy but due to envy.

### Affiliative rats emitted 22-kHz calls of short duration and high frequency during the inequality conditions

We then investigated the acoustic characteristics of calls emitted during social situations compared with those during air puff stress. Air puffs were applied to a separate series of non-stroked female rats. USVs during air puff stress had a low peak frequency, particularly below 30-kHz (Fig. 3a, c). On the other hand, the frequency of 22-kHz calls emitted during social isolation (condition 1) or social inequality conditions (conditions 4 and 5) was higher than the frequency in the air-puff condition (Fig. 3b, d, f, g). In condition 2, there was a large number of calls around 50-kHz (Fig. 3e). For analysis of the average frequency, duration, and amplitude, we combined all USVs between the 18 kHz band and 32 kHz band. The mean frequencies of 22-kHz calls (range of 18-32 kHz) in social conditions 1 ((28.05 ± 0.37 (mean ± SEM, n = 12)), 2 (28.4 ± 0.94, n = 12), 4 (27.63 ± 0.79, n = 12), and 5 (28.79 ± 0.57, n = 12) were higher than the frequency in the air-puff condition (22.33 ± 0.46, n = 9) (χ2 = 20.995, *df* = 4, *P* < 0.001, Kruskal-Wallis test, air-puff condition versus condition 1, *P* = 0.026; air puff condition versus condition 2, *P* < 0.001; air puff condition versus condition 4, *P* = 0.019; air-puff condition versus condition 5, *P* = 0.001, post-hoc Holm’s test) (Fig. 3h). The mean durations of 22-kHz calls in conditions 1 (0.065 ± 0.01 (mean ± SEM, n = 12), 2 (0.047 ± 0.005, n = 12), 4 (0.059 ± 0.01, n = 12), and 5 (0.075 ± 0.001, n = 12) were shorter than the mean duration in the air-puff condition (0.722 ± 0.098, n = 9) (χ2 = 18.140, *df* = 4, *P* = 0.001, Kruskal-Wallis test, air-puff condition versus condition 1, *P* = 0.032; air-puff condition versus condition 2, *P* = 0.001; air-puff condition versus condition 4, *P* = 0.01, post-hoc Holm’s test) (Fig. 3i). There were no significant differences among the conditions in the mean amplitudes of calls (frequency range of 18-32 kHz) (Fig. 3j).

**Fig.3:**
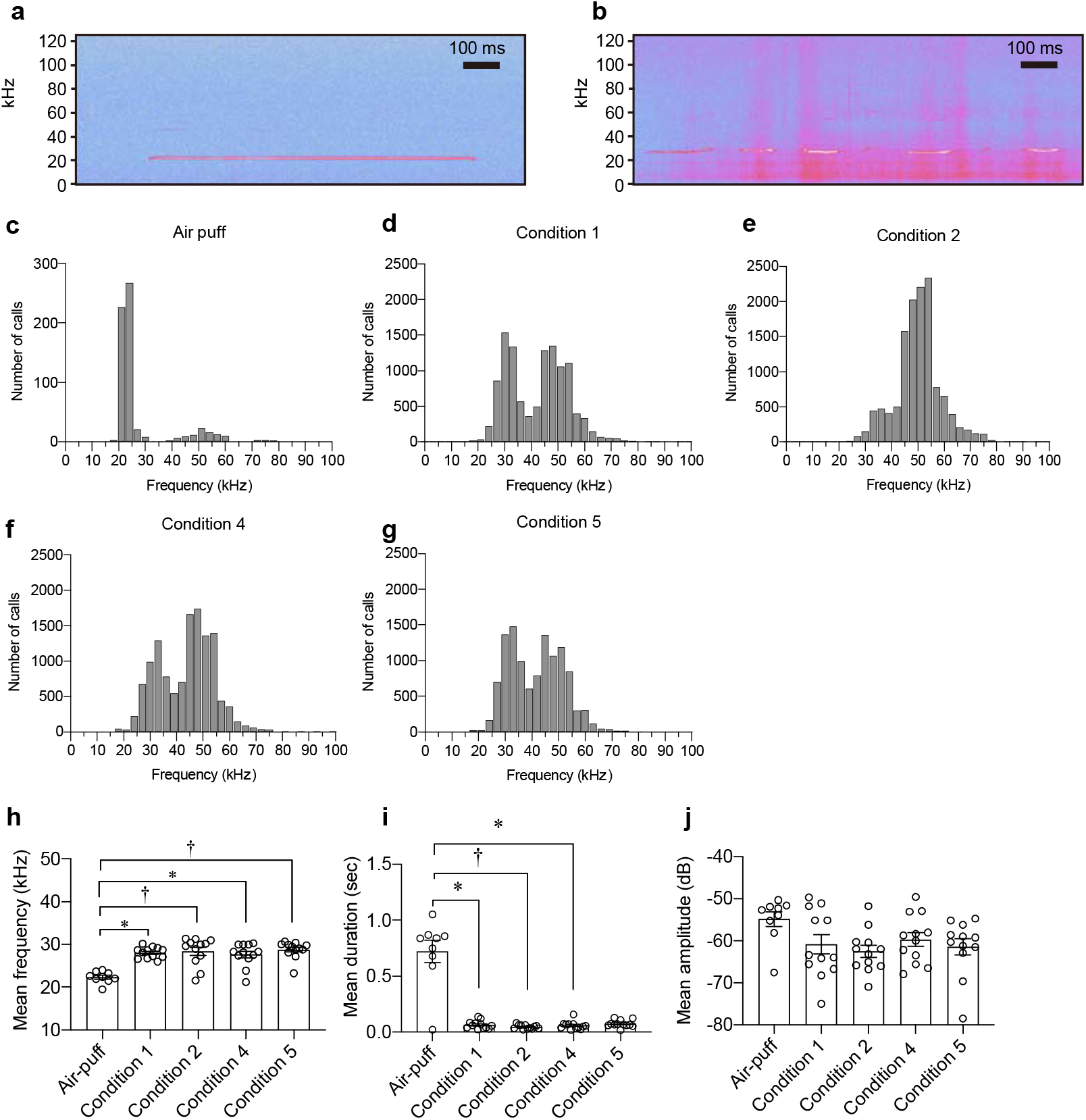
Affiliative rats emitted 22-kHz calls of short duration and hight frequency during the social inequality conditions. Spectrograms of 22-kHz calls during the air-puff condition (a) and during an inequality condition (condition 4) (b). Histogram of the number of USVs of LEW female rats (N = 9) in the air-puff condition (c). USVs of affiliative group rats (N = 12) in condition 1 (d), condition 2 (e), condition 4 (f), and condition 5 (g). Mean frequencies (h), durations (i), and amplitudes (j) of calls (frequency range of 18-32 kHz) in the air-puff condition and conditions 1, 2, 4, and 5. †, *P* < 0.01, *, *P* < 0.05. Error bars denote standard error of the mean.

### An oxytocin receptor antagonist did not affect the social preference and emission of USVs during social inequality conditions

Our previous studies suggested that the affiliative response toward a human or gentle stroking stimuli was correlated with the neural activities of oxytocin neurons in the caudal PVN ^8^. Thus, we examined the role of the oxytocin receptor in social preference toward an experimenter and emission of USVs in inequality conditions. Thirty min after intraperitoneal injection of an oxytocin receptor antagonist (OTA) or saline, a social preference test was conducted. Repeated measures two-way ANOVA revealed a significant effect of experimenter in times spent in hand areas [F (1, 10) = 19.431, *P* = 0.001]. However, there was no significant effect of drug and interaction of the experimenter and drug (Fig. 4a). Rats in the OTA-injected and saline-injected groups showed significantly higher preference toward a familiar experimenter than an unfamiliar experimenter. The results indicate that the oxytocin receptor did not play a crucial role in expression of social preference toward a familiar experimenter. In addition, there was no significant difference in the numbers of 22-kHz and 50-kHz calls during the social inequality condition in which other rats received stroking stimuli from a familiar experimenter between the antagonist-injected rats and saline-injected rats (Wilcoxon signed-rank test) (Fig. 4b-d). This result suggests that the oxytocin receptor was not necessary for emission of USVs during an inequality condition.

**Fig.4:**
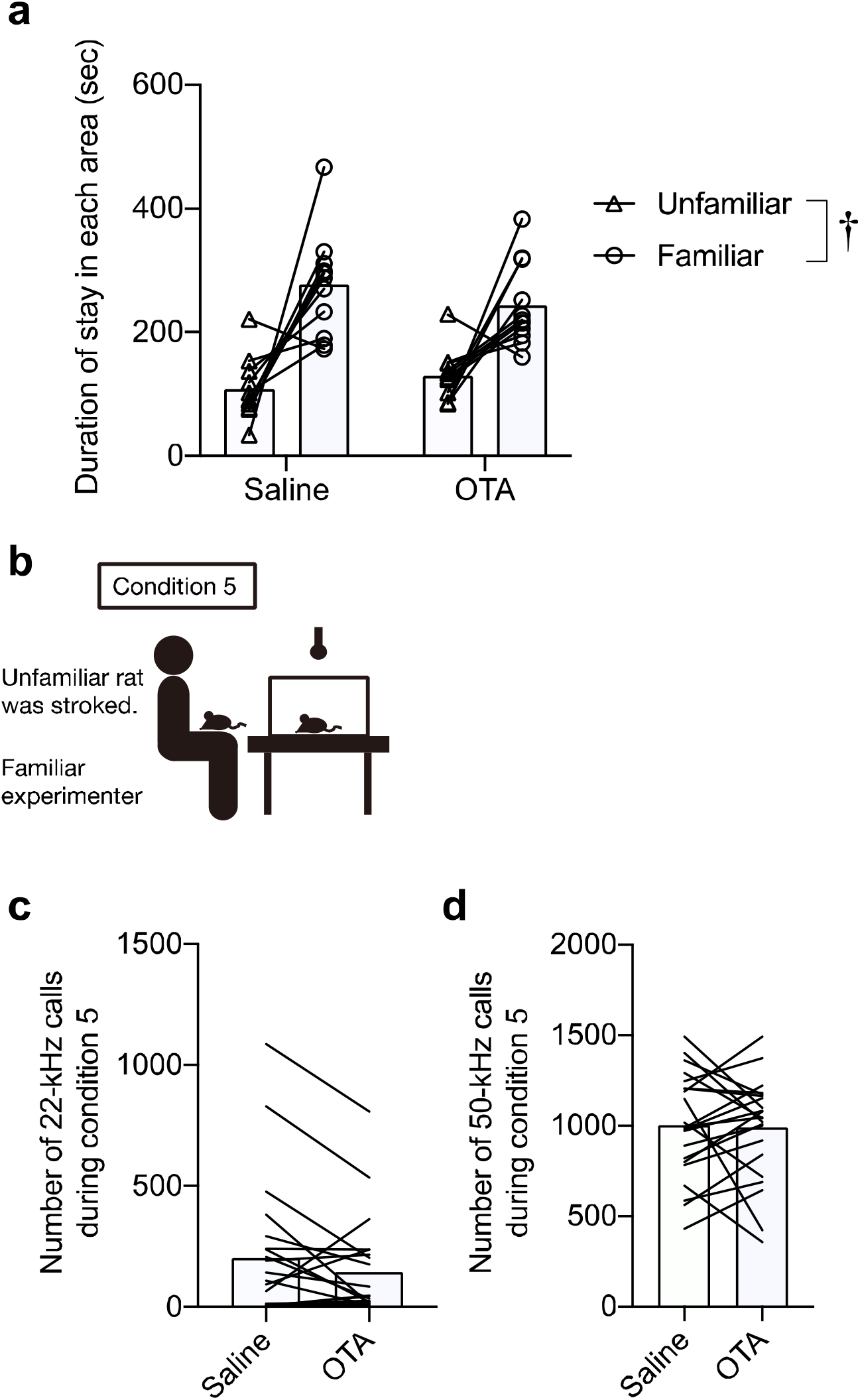
Oxytocin antagonist did not affect the social preference and emission of USVs during inequality conditions. (a) Durations of staying in the familiar experimenter’s hand area and unfamiliar experimenter’s hand area in the social preference test after administration of an oxytocin receptor antagonist (OTA) or saline. (b) Illustration of experimental procedures. Thirty min after administration of OTA or saline, observer rat’s USVs were recorded when a familiar experimenter stroked unfamiliar rats in front of the observer’s cage. Numbers of 22-kHz (c) and 50-kHz (d) calls. For both 22-kHz and 50-kHz calls. †, *P* < 0.01.

## Discussion

In the present study, we revealed that rats that repeatedly received affiliative touches during the developmental period showed a specific social preference toward a familiar experimenter who repeatedly interacted with the rats. In addition, we revealed that rats emitted a larger number of 50-kHz calls when the familiar experimenter gently stroked the rats than the number of calls when an unfamiliar experimenter stroked them. These rats emitted 22-kHz calls during social isolation, but the number of 22-kHz calls was dramatically decreased when a familiar experimenter stayed in front of the rats but not when an unfamiliar experimenter was present. These results revealed that rats could discriminate humans and that the presence of familiar humans decreased negative emotion during isolation and increased positive emotion during stroking stimuli. These rats appeared to have a social bond with the familiar experimenter. A social bond has been defined by behavioral or physiological responses: preference for the attachment figure, maintenance of proximity or voluntary contact with it, increased stress responses when separated from it and decreased stress responses when reunited^15^. Our previous study revealed that repeated stroking stimuli induce the following behavior of the rat toward the familair human. Considering the behavioral responsiveness of affiliative rats in our previous study and the present study, it is thought that repeated stroking during the developmental period can result in social bonding between humans and rats.

We showed that affiliative rats emitted 22-kHz calls during inequality conditions. Rats emitted a larger number of 22-kHz calls during isolation and an inequality condition but not when a familiar experimenter was present in front of them, suggesting that rats discriminated social conditions and showed negative emotion in socially isolated and inequality conditions. The aversive property of inequitable outcomes has been demonstrated in several non-human primates ^16–18^ and other social species including domestic dogs ^3,4^, corvids^5^, and rodents ^6,7^. It is possible that inequality aversion and sociality, particularly cooperation, may have coevolved ^19^. Rats are a highly social species and form a hierarchically structured, well-organized social group. Several studies have suggested that rats show coordinated cooperative actions ^20^, reciprocity ^21^, empathy and prosocial behavior ^22,23^. In addition, Kashtelyan and colleagues reported that rats emitted 22-kHz calls during observation of conspecific reward, food delivery^24^. However, several studies have failed to demonstrate inequality aversion in social species ^25–27^. It has been argued that costly refusals of unfair offers, as demonstrated in several primate studies, may merely reflect nonsocial motives such as frustration effects and/or violated expectations. However, frustration, expectation violations or other nonsocial motives are unlikely to fully explain the emission of 22-kHz calls in our study. Rats in the affiliative group emitted a larger number of 22-kHz calls both during social isolation and in an inequality condition but not in the presence of the familiar experimenter in front of them (condition 2). If experimental environments including the existence of the experimenter induced anticipation of social reward, rats should have emitted 22-kHz calls during condition 2 in which the familiar experimenter stayed in front of the rats. We conclude that nonsocial motives such as betray of expectation are unlikely to fully explain the distress calls by rats in our study.

Interestingly, 22-kHz calls by rats in the affiliative group during social isolation and in inequality conditions had different acoustic properties from those emitted in the air puff condition. These results suggest that rats exhibit different distress calls depending on conditions. Several studies revealed that young rats (4 weeks of age) emitted 22-kHz calls with a higher frequency and shorter duration than did adult rats (11 or 12 weeks of age) ^28,29^. In this study, the age of rats in the air-puff condition was 14 weeks, and the ages of rats in conditions 1 – 4 were 12 – 16 weeks. Thus, the difference in acoustic characteristics of 22-kHz calls was not due to age^10^. Since rats produce USVs with different acoustic characteristics than those in an air-puff condition, they might exhibit distinct unpleasant emotions of social isolation and inequality that are different from those of invasive physical stimuli. Brudzynski reported that 22-kHz calls can be subdivided into long and short forms of calls ^30^ and that a short duration of 22-kHz calls was observed during withdrawal from cocaine^31^. Previous studies reported that a negative emotion associated with an internal discontent is expressed by short calls ^11, 32^. From these studies, shorter (0.05-0.07-sec duration) and higher frequency (28 kHz) 22-kHz calls in our study may be characteristics of ultrasonic vocalization emitted during negative emotion associated with social relationships. Further examination of USVs and their effects on behaviors could shed light on the physiological meaning of USVs in social isolation and inequity conditions.

Studies concerning inequity aversion in animals have focused on behavioral responses, and the neural mechanisms underlying inequality aversion remain to be clarified. Several studies suggested that dopamine, serotonin, and oxytocin modulate the egalitarian and trusting behaviors in human^33–35^. Oxytocin has been shown to modulate affiliative social behaviors in animals^36^. In addition, oxytocin has been reported to modulate the responses to inequity in dogs^37^. From those studies, we expected that administration of an oxytocin receptor antagonist would affect the response to an inequality condition in rats. However, contrary to our expectation, the oxytocin receptor antagonist had no significant effect on the emission of 22-kHz USVs in an inequality condition as well as social preference behavior. It is possible that oxytocin contributes mainly to the building process of social relationships, not to the expression of affiliation-related responses. Further research is needed to clarify the neural mechanisms of inequality aversion.

The results of the present study suggested that rats express negative emotion in social inequality conditions. This study provides an experimental animal model to investigate molecular and neural mechanisms underlying affiliative relationships and inequality aversion.

## Lead contact and availability of materials

Further information and requests for resources should be directed to Shota Okabe or Tatsushi Onaka. This study did not generate new unique reagents.

## Data and code availability

The datasets supporting the current study have not been deposited in a public repository but are available from the corresponding author on request. This study did not generate code.

## Experimental model and subject details

### Ethical approval

Animal experiments were carried out after receiving approval from the Animal Experiment Committee of Jichi Medical University and were in accordance with the Institutional Regulations for Animal Experiments and Related Activities in Academic Research Institutions under the Jurisdiction of the Ministry of Education, Culture, Sports, Science and Technology.

### Animals

Ninety-seven female rats of the Lewis strain (LEW/ CrlCrlj, Charles River Laboratories Japan, Inc., Kanagawa, Japan) were used in the present study. The rats were obtained from an animal supplier at 3 weeks of age. They were housed in pairs under a 12: 12h light/dark cycle (lights on at 7:30 am) at 22 ± 2°C and 55 ± 15% relative humidity. Food and water were available *ad libitum*.

### General experimental design for preparing the animals

In the affiliative group, rats (N = 80) were kept in pairs and received affiliative touches 3 to 5 times a week from an experimenter (S.O.) for 4 weeks between 3 and 6 weeks of age. Affiliative touches consisted of affiliative interactions such as stroking and tickling. Stroking stimuli were applied by an experimenter (S.O.) with the hand of the experimenter over the back of each animal at a speed of 5-10 cm/sec. Tickling stimuli consisted of three phases: dorsal tickling, flip (rapidly overturned), and ventral tickling (vigorous tickling on the ventral trunk while the animal was pushed on the floor). Each phase lasted approximately 5-10 sec and the duration of one tickling stimulus was15-30 sec. Tickling was repeated at intervals of 15 sec. A pair of rats received stroking stimuli at the same time. The experimenter wore cotton gloves and a lab coat during interaction with the rats. Rats in the control group (N = 8) were housed in pairs and did not receive affiliative touches until the experiments. In addition, another group of female rats at 14 weeks of age was used for recording USVs during the air puff condition (N = 9).

USVs in the affiliative group were recorded during a 5-min stroking session once each week during the developmental period (3-6 weeks of age). Rats were individually placed on the experimenter’s (S.O.) lap and each rat was given massage-like stroking stimuli from the experimenter while on the experimenter’s lap. The numbers of 50-kHz calls (frequency between 35 and 100 kHz) and 22-kHz calls (frequency between 18 and 32 kHz) were automatically counted using our custom-written MATLAB codes as reported previously ^38^. At the age of 6 weeks, we selected rats for further experiments according to the total number of 50-kHz calls recorded at 3, 4, 5, and 6 weeks of age in response to stroking stimuli. The top 34 out of the 80 rats in the affiliative group were selected. These selected rats in the affiliative group received affiliative touches 3 to 5 times a week for another 6 weeks between 7 and 12 weeks of age. Thus, rats in the affiliative group received affiliative touches for 10 weeks in total between 3 and 12 weeks of age. Twelve rats in the affiliative group were used in experiments for which results are shown in Fig. 1a-g, 2a-c, and 3a-j, and 22 rats in the affiliative group were used in experiments for which results are shown in Fig. 1h-m, 2d-f, and 4.

### General protocols for recording USVs

USVs were recorded by the use of a microphone (Type 4158 N, Aco, Tokyo, Japan: CM16, Avisoft, Berlin, Germany) and an A/D converter (Ultra-SoundGate, Avisoft Bioacoustics, Berlin, Germany; SpectoLibellus 2D, Katou Acoustics Consultant Office, Kanagawa, Japan). The microphone was placed approximately 20 cm from the rat (see section “Recording of USVs during each condition for detailed description).

### Recording of USVs during each condition

Condition 1: Each rat was placed and kept in a clean cage that was the same as its home cage. USVs were recorded for 15 min.

Condition 2: Each rat was kept in a clean cage as in condition 1 and a familiar experimenter (S.O.) sat down in front of the cage (at a distance of approximately 60 cm from the cage). USVs were recorded for 15 min. The experimenter remained seated and tried not to move.

Condition 3: Each rat was kept in a clean cage as in condition 1 and an unfamiliar experimenter sat down in front of the cage (at a distance of approximately 60 cm from the cage). USVs were recorded for 15 min. The experimenter remained seated and tried not to move.

Condition 4: Each rat was kept in a clean cage as in condition 1 and a familiar experimenter stroked its cage mate in front of the cage at a distance of approximately 60 cm from the cage. USVs were recorded for 15 min.

Condition 5: Each rat was kept in a clean cage as in condition 1 and a familiar experimenter (S. O.) stroked another rat (demonstrator) that was unfamiliar to the subject rat in front of the cage at a distance of approximately 60 cm from the cage. USVs were recorded for 15 min.

Condition 6: Each rat was kept in a clean cage as in condition 1 and an unfamiliar experimenter stroked another rat (demonstrator) that was unfamiliar to the subject rat in front of the cage at a distance of approximately 60 cm from the cage. USVs were recorded for 15 min.

Air puff condition: To elicit distress 22-kHz calls, another group of rats were individually transferred to an experimental cage and habituated to the cage for 5 min. Then each rat received air-puff stimuli (0.3 MPa) with an inter-stimulus interval of 2 s to the nape from a distance of approximately 5-10 cm. Immediately after 30 air-puff stimuli had been delivered, USVs were recorded for 5 min. The data used in our previous study concerning the establishment of algorithms of software programs (USVSEG) for automatic analysis of USVs^38,39^ were used.

The number of 50-kHz calls decreased and the number of 22-kHz calls increased continuously during the first 10 min in affiliative rats in the isolation condition (condition 1) (Supplementary Fig. 1a, b). In contrast to rats in the affiliative group, rats in the control group emitted a smaller number of USVs and there were no significant changes in the number of calls (Supplementary Fig. 1c, d). Thus, we used calls for the first 10 min for analysis (condition 1 to 6).

### Social preference test

A social preference test was used for assessment of preference to the experimenter (S.O.) ‘s hand in comparison with the unfamiliar experimenter’s hand. The experimenter (S.O.) ‘s hand and the unfamiliar experimenter’s hand were placed at diagonally opposite corners of a test box (W60 × D60 × H40 cm). Each animal was placed in the test box at a corner different from the corners where hands were placed. The time spent staying in a square (20 × 20 cm) of the hand area was measured automatically during a 10-min test period using a program (Time OFCR1, O’Hara & CO., LTD, Tokyo, Japan).

### Intraperitoneal administration

The non-peptidergic oxytocin receptor antagonist L-368,899 (R&D Systems, MN, USA) was dissolved in saline (0.9%) at a concentration of 0.2 mg/ ml and administered to rats via intraperitoneal (IP) injection at a dose of 1 mg/ kg body weight.

### Experimental schedule

USVs of selected rats in the affiliative group in the first series of sessions (N = 12) and rats in the control group (N = 8) were first recorded in condition 1. On days 2 and 3, USVs of the rats were recorded in condition 2. On days 4 and 5, USVs of the rats were recorded in condition 4. Rats were housed in pairs and one of the pair was assigned to be the demonstrator and the other was assigned to be the observer in condition 4. The experiments for condition 4 were conducted over a period of 2 days, and the order of the roles of observer and demonstrator was counterbalanced; the observer rats on day 4 were assigned to be the demonstrator rats on day 5 and the demonstrator rats on day 4 were assigned to the observer rats on day 5. On day 6, rats were examined for social preference toward experimenters in the social preference test. On the last day, USVs of rats were recorded in condition 5. All of the tests were performed for all rats.

Selected rats in the affiliative group in the second series of sessions (N = 22) were first examined for social preference toward experimenters in the social preference test on day 1. On days 2 and 3, USVs of rats were recorded in conditions 5 and 6. The experiments were conducted over a period of 2 days, and the order of experimenters was counterbalanced. On days 4 and 5, USVs of half of the rats (N = 11) were recorded in condition 5 at 30 min after IP administration of an oxytocin receptor antagonist or saline. The experiments were conducted over 2 days, and the order of drugs was counterbalanced. On days 6 and 8, the same rats (N = 11) were tested for social preference 30 min after IP administration of an oxytocin receptor antagonist or saline. The experiments were conducted over a period of 2 days, and the order of drugs was counterbalanced. On days 7 and 9, USVs of the other half of the rats (N = 11) were recorded in condition 5 at 30 min after IP administration of an oxytocin receptor antagonist or saline. The experiments were conducted over a period of 2 days, and the order of drugs was counterbalanced. Thus, the total number of animals in condition 5 with drug administration was 22, and the total number of animals in the social preference test with drug administration was 11. On day 10, USVs of all rats were recorded in condition 1. On days 11 and 12, USVs of rats were recorded in conditions 2 and 3. The experiments were conducted over a period of 2 days, and the order of conditions was counterbalanced. The number of rats was 20 due to equipment trouble. On days 13 and 14, USVs of rats were recorded during stroking stimuli by a familiar experimenter or unfamiliar experimenter. The unfamiliar experimenter learned the stroking procedures before experiments. IP administration was conducted by an experimenter blind to treatments of the animals and the experimenter of behavioral tests was blind to treatments of IP administration.

### Quantification and statistical analysis

Statistical analysis was performed using free software, HAD ^39^. Developmental changes in the numbers of USVs in the affiliative group (Fig. 1d, e), time course of the number of USVs for 15 min in condition 1 (Supplementary Fig. 1a-d), and number of USVs during conditions 1, 2, and 3 (Fig. 1l, m) were analyzed by repeated-measures one-way ANOVA followed by post-hoc Holm’s test. Results of the social preference test shown in Figures 1g and 4a were analyzed using repeated two-way ANOVA (group × experimenter, drugs × experimenter) followed by post-hoc Holm’s test. The numbers of USVs during conditions 1, 2, 4, and 5 (Fig. 2b, c) were also analyzed using repeated two-way ANOVA (group × condition). Acoustic characteristics in the air-puff condition and conditions 1, 2, 4, and 5 were analyzed by the Kruskal-Wallis test followed by post-hoc Holm’s test (Fig. 3h-j). Differences in the numbers of USVs between a familiar experimenter and unfamiliar experimenter (Fig. 1l, j) and between conditions 5 and 6 (Fig. 2e, f) were analyzed by Wilcoxon’s signed-rank test. The number of USVs after administering drugs was also analyzed by Wilcoxon’s signed-rank test (Fig. 4c, d). *P* < 0.05 was considered statistically significant.

## Acknowledgements

This work was financially supported by the Japan Society for the Promotion of Science, Grants-in Aid for Scientific Research from the Ministry of Education, Culture, Sports, Science, and Technology of Japan (#15K18978, #17H05956 and #20K14260 to S.O.; #17K08573 and #20K07278 to Y.T.; #17K08574 and #20K07264 to M.Y.; #17H04026, #19K22475 and #20H03419 to T.O.). The funding agencies had no role in the design of the study, data acquisition and analysis, or manuscript preparation.

## Author contributions

S.O. and Y.T. performed the experiments and analyzed the behavioral data and Y.T., M.Y., and T.O. supervised the study. All authors wrote and revised the manuscript and approved the final manuscript.

