## Supplementary Information for "Rats emit unique distress calls in social inequality conditions"

**Supplementary Movie 1: Observer rats emitted distress calls when demonstrator rats received stroking stimuli in front of them.**

**Supplementary Movie 2: Rats in the affiliative group, but not rats in the control group, emitted distress calls during an inequality condition.**

**Supplementary Fig. 1: Changes in the number of USVs over time.**

**Supplementary Movie 1: Observer rats emitted distress calls when demonstrator rats received stroking stimuli in front of them.**

　The experimenter stroked demonstrator rat in front of the cage containing observer rat. The audio signal provides ultrasonic vocalizations that have been transposed to the human audible frequency range.

**Supplementary Movie 2: Rats in the affiliative group, but not rats in the control group, emitted distress calls during an inequality condition.**

　Distress calls of rats in the affiliative and control groups when the experimenter stroked a demonstrator rat in front of the cage containing an observer rat. The audio signal provides ultrasonic vocalizations that have been transposed to the human audible frequency range.

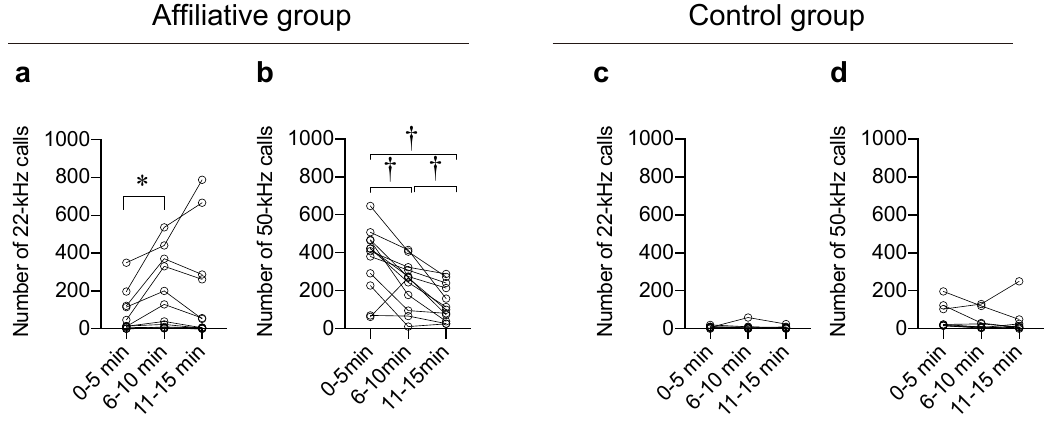

**Supplementary Fig. 1. Changes in the number of USVs over time.**

Time courses of the numbers of vocalizations in the affiliative group (a, 22-kHz; b, 50-kHz) and control group (c, 22-kHz; d, 50-kHz) during condition 1. In the number of 22-kHz calls in the affiliative group, repeated measures one-way ANOVA revealed a significant effect of time [F (2, 22) = 3.952, *P* = 0.046]. The number of 22-kHz calls in 6-10 min was significantly larger than that in 0-5 min (a, *P* = 0.048, post-hoc Holm's test). In the number of 50-kHz calls in the affiliative group, repeated measures one-way ANOVA revealed a significant effect of time [F (2, 22) = 20.726, *P* < 0.001]. The numbers of 50-kHz call in 6-10 min and 11-15 min were significantly smaller than the number of 50-kHz calls in 0-5 min (b, 0-5 min versus 6-10 min, *P* = 0.007; 0-5 min versus 11-15 min, *P* = 0.001, post-hoc Holm’s test). The number of 50-kHz calls in 11-15 min was significantly smaller than that in 6-10 min (*P* = 0.002, post-hoc Holm's test). There were no changes in the number of 22-kHz (c) and 50-kHz (d) calls in the control group. *, *P* < 0.05, †, *P* < 0.01.
